# Macrodomain Mac1 of SARS-CoV-2 Nonstructural Protein 3 Hydrolyzes Diverse ADP-ribosylated Substrates

**DOI:** 10.1101/2023.02.07.527501

**Authors:** Chanbora Chea, Duck-Yeon Lee, Jiro Kato, Hiroko Ishiwata-Endo, Joel Moss

## Abstract

Severe acute respiratory syndrome coronavirus 2 (SARS-CoV-2) is responsible for a global pandemic that resulted in more than 6-million deaths worldwide. The virus encodes several non-structural proteins (Nsps) that contain elements capable of disrupting cellular processes. Among these Nsp proteins, Nsp3 contains macrodomains, e.g., Mac1, Mac2, Mac3, with potential effects on host cells. Mac1 has been shown to increase SARS-CoV-2 virulence and disrupt ADP-ribosylation pathways in mammalian cells. ADP-ribosylation results from the transfer of the ADP-ribose moiety of NAD^+^ to various acceptors, e.g., proteins, DNA, RNA, contributing on a cell’s biological processes. ADP-ribosylation is the mechanism of action of bacterial toxins, e.g., Pseudomonas toxins, diphtheria toxin that disrupt protein biosynthetic and signaling pathways. On the other hand, some viral macrodomains cleavage ADP-ribose-acceptor bond, generating free ADP-ribose. By this reaction, the macrodomain-containing proteins interfere ADP-ribose homeostasis in host cells. Here, we examined potential hydrolytic activities of SARS-CoV-2 Mac1, 2, and 3 on substrates containing ADP-ribose. Mac1 cleaved α-NAD^+^, but not β-NAD^+^, consistent with stereospecificity at the C-1” bond. In contrast to ARH1 and ARH3, Mac1 did not require Mg^2+^ for optimal activity. Mac1 also hydrolyzed *O*-acetyl-ADP-ribose and ADP-ribose-1”-phosphat, but not Mac2 and Mac3. However, Mac1 did not cleave α-ADP-ribose-(arginine) and ADP-ribose-(serine)-histone H3 peptide, suggesting that Mac1 hydrolyzes ADP-ribose attached to O- and N-linked functional groups, with specificity at the catalytic site in the ADP-ribose moiety. We conclude that SARS-CoV-2 Mac1 may exert anti-viral activity by reversing host-mediated ADP-ribosylation. New insights on Nsp3 activities may shed light on potential SARS-CoV-2 therapeutic targets.

**IMPORTANCE:** SARS-CoV-2, the virus responsible for COVID-19, encodes 3 macrodomain-containing proteins, e.g., Mac1, Mac2, Mac3, within non-structural proteins 3 (Nsp3). Mac1 was shown previously to hydrolyze ADP-ribose-phosphate. Inactivation of Mac1 reduced viral proliferation. Here we report that Mac1, but not Mac2 and Mac3, has multiple activities, i.e., Mac1 hydrolyzed. α-NAD^+^ and *O*-acetyl-ADP-ribose. However, Mac1 did not hydrolyze β-NAD^+^, ADP-ribose-serine on a histone 3 peptide (aa1-21), and ADP-ribose-arginine, exhibiting substrate selectivity. These data suggest that Mac1 may have multi-function as a α-NAD^+^ consumer for viral replication and a disruptor of host-mediated ADP-ribosylation pathways. Understanding Mac1’s mechanisms of action is important to provide possible therapeutic targets for COVID-19.

## INTRODUCTION

COVID-19, a severe acute respiratory syndrome mediated by SARS-CoV-2 generated in a worldwide public health emergency, with greater than 6.7 million deaths according to World Health Organization (WHO; https://covid19.who.int). This virus has emerged in 2019 as the third coronavirus (CoV) mediated disease in the past two decades (1). In 2002-2003, severe acute respiratory syndrome-related coronavirus (SARS-CoV) was reported in about 8000 infected cases with approximately 10% deaths, followed by an arrival of Middle East respiratory syndrome coronavirus (MERS-CoV) in 2012 with about 2,500 infected people with a 35% mortality rate (2).

ADP-ribosylation, a post-translational modification (PTM), is catalyzed by ADP-ribosyltransferases (ARTs) including poly(ADP-ribosyl)polymerases (PARPs) that transfer ADP-ribose (ADPr) from nicotinamide adenine dinucleotide (NAD^+^) to target substrates via *N*-, *O*-, or *S*-glycosidic linkages at the 1” position of the nicotinamide ribose via an SN2-like reaction mechanism (3–5). A monomeric or polymeric reaction product, resulting in, respectively, mono ADP-ribose (MAR) or a poly(ADP-ribose) (PAR) suppresses viral replication (6), i.e., PARPs inhibit virus replication (7). Supporting their anti-viral activities, some PARPs are activated by interferon (IFN), suggesting that ADP-ribosylation is responsible for IFN-dependent innate immunity upon virus infection (8, 9).

Hydrolytic enzymes, <u>e.g., ADP-ribosylhydrolases (ARHs), poly (ADP-ribose) glycohydrolase (PARG), and macrodomain-containing proteins,</u> have been shown to control ADP ribosylation by cleaving ADP ribose from the target protein, DNA, and RNA, <u>which involve</u> signaling pathways and biological functions (10, 11).

SARS-CoV-2 consists of a large RNA genome encoding 29 proteins, including structural, nonstructural, and accessory proteins, which are involved in viral functions and virulence (12). Among the 16 nonstructural proteins is which nonstructural protein 3 (Nsp3), a 200 kDa multi-domain protein (13). The Nsp3 proteins possess three tandem macroD-like domains (e.g., Mac1, Mac2, Mac3) (14). SARS-CoV-2 Mac1, contains three-layer α/β/α fold and binds to mono-ADP-ribose (MAR) (15). However, Mac2 and Mac3 failed to interact with ADP-ribose, but do bind to nucleic acids (16, 17). SARS-CoV-2 Mac1 was responsible for viral replication in some cell lines *in vitro* (18, 19) and promotes immune evasion *in vivo* (13, 20, 21).

SARS-CoV-2 Mac1 is believed to be a mono (ADP-ribosyl) hydrolase, removing single ADP-ribose from ADP-ribosylated protein in cells to promote viral replication, and inhibit host immune response including interferon (IFN) and interferon stimulated genes (ISGs) (22–24). Thus, the host immune response fails to produce potent IFN responses, resulting in induction of chemokines (25). Likewise, the activation of ADP-ribosylation by PARP9/DTX3L heterodimer following IFN stimulation was reversed by ectopic expression of the SARS-CoV-2 Nsp3 macrodomain in human cells (26).

As described above, enzymatic activity of SARS-CoV-2 Mac1 is essential for virus infection. Identification of substrates Mac1 in host cells may help define its cellular functions and provide alternative therapeutic targets.

## RESULTS

### SARS-CoV-2 Mac1 hydrolyzed α-NAD^+^ but not α-ADP-ribose-arginine or β-NAD^+^

To determine whether SARS-CoV-2 Mac1 protein hydrolyzes N-linked ADP-ribosylated substrates, Mac1 was incubated with various substrates including α-NAD^+^, β-NAD^+^, and α-ADPr-arginine. The results showed that SARS-CoV-2 Mac1 cleaved α-NAD^+^ to release the ADP ribose and nicotinamide (Figure 1, IB) with the reaction products confirmed by MS analysis. The SARS-CoV-2 Mac1 showed about 2.3-times lower α-NADase activity than that of ARH3 (positive control). In contrast, β-NAD^+^ or α-ADPr-arginine hydrolysis were not observed (data not shown). These results indicated the stereospecific hydrolysis of α-NAD^+^ by macrodomain Mac1 of SARS-CoV-2 as an α-ADPr-acceptor hydrolase.

**Figure 1:**
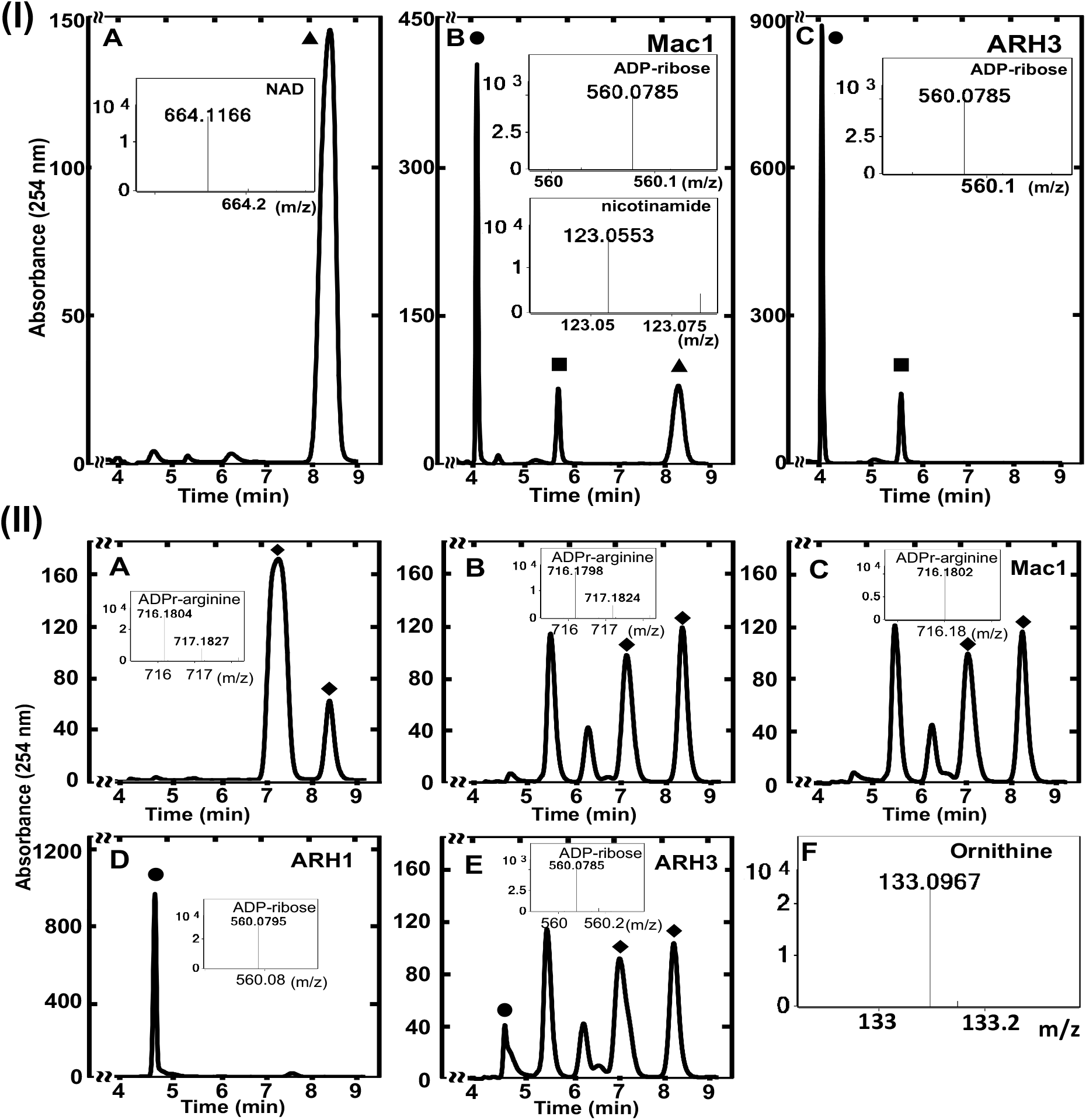
The SARS-CoV-2 Mac1 hydrolyzed α-NAD^+^ but not α-ADP-ribose arginine. Proteins were incubated overnight at 37°C, in 200μl of reaction mix, with substrates in 50 mM potassium phosphate pH 7.0, 10 mM MgCl_2_, 5 mM DTT, and 22.5 μM ovalbumin. (I) Reaction products from SARS-CoV-2 Mac1 with α-NAD^+^. (A) α-NAD^+^ (20 μM), (B) Mac1 (65 nM), (C) ARH3 (65 nM). (II) Hydrolysis of ADP-ribose-arginine by SARS-CoV2 Mac1. (A) ADP-ribose-arginine (7 nmols), (B) ADP-ribose-arginine (37°C, overnight), (C) SARS-CoV-2 Mac1 (65 nM), (D) human ARH1 (65 nM), (E) human ARH3 (650 nM), (F) Ornithine released. Incubated samples were then separated by RP-HPLC and monitored at 254 nm as described in Materials and Methods. Purified peaks were confirmed by mass spectrometry. Samples were done in duplicate, and data represented a single separation. (▲) α-NAD^+^, (●) ADP-ribose, (◼) nicotinamide, and (◆) ADP-ribose arginine.

We found that neither Mac2 nor Mac3 hydrolyzed α-NAD^+^, β-NAD^+^ or α-ADPr-arginine (data not shown). α-ADPr-arginine was hydrolyzed by ARH1 (positive control) with significantly less activity with ARH3 (positive control) (Figure 1, IID, IIE). The data showed that SARS-CoV-2 Mac1 did not hydrolyze β-NAD^+^, consistent with it being stereospecific at the C-1” bond.

### ADPr-1”-phosphate and *O*-Acetyl-ADPr were substrates of SARS-CoV-2 Mac1 but not ADP-ribose-serine (ADPr-Ser)

*O*-linked functional groups including *O*-Acetyl-ADPr were reported to deacetylate by a family of human macrodomain proteins, MacroD1 and MacroD2 (27), and ADPr-phosphate was dephosphorylated by Af1521 (15).

We first examined the cleavage activity of SARS-CoV-2 Mac1 on *O*-Acetyl-ADPr. Using incubated samples of enzyme with *O*-Acetyl-ADPr as a substate, the products were separated by RP-HPLC as described in Materials and Methods. We found that Mac1 deacetylated *O*-Acetyl-ADPr and generated ADPr similar to that was seen with ARH3 as positive control; the products were confirmed by mass spectrometry (Figure 2, I). Mac2 and Mac3 did not have deacetylate *O*-Acetyl-ADPr (data not shown). In the absence of Mg^2+^ from the assay reaction mix, we found that SARS-CoV-2 Mac1 hydrolyzed both α-NAD^+^ to release ADPr and nicotinamide and *O*-Acetyl-ADPr to release ADPr.

**Figure 2:**
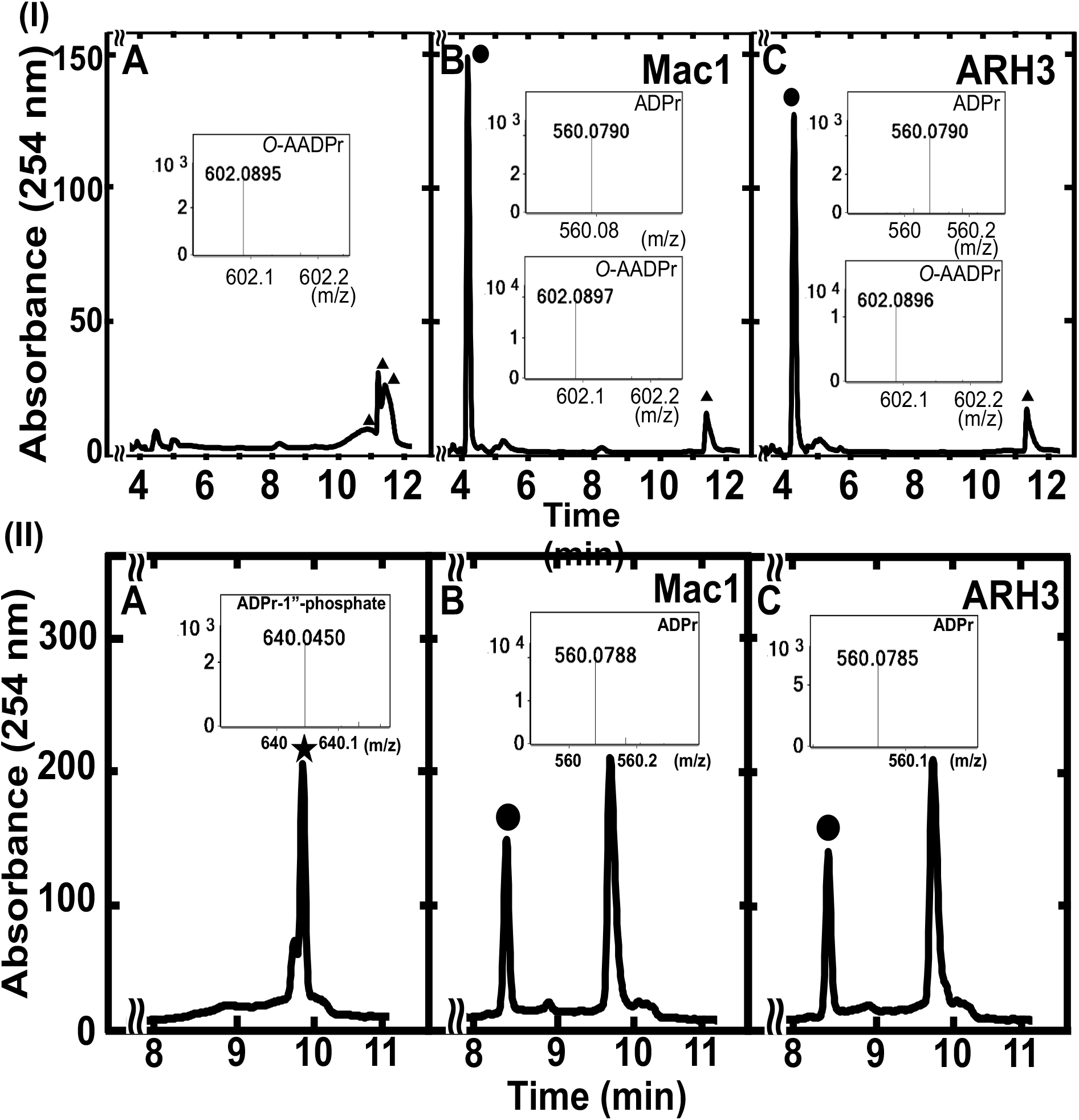
Hydrolysis of *O*-acetyl-ADP-ribose (*O*AADPr)and ADPr-1”-phosphate by Mac1. (I) The SARS-CoV-2 Mac1 hydrolyzed *O*AADPr. Mac1 was incubated at 25°C for 1 h, in 200 μl of reaction mixture, with 50 mM potassium phosphate pH 7.0, 10 mM MgCl_2_, and 22.5 μM ovalbumin. (A) (▲) *O*AADPr (20 μM), (B) Mac1 (65 nM), (C) ARH3 (65 nM). (II) Mac1 was incubated overnight at 37°C, in a 200 μl of mixture, with ADPr-1”-phosphate, in 50 mM Tris-HCl (pH 7.5), 10 mM MgCl_2_, 5 mM DTT, and 22.5 μM ovalbumin. (A) (★) ADP-ribose-1”-phosphate (6 μM), (B) Mac1 (650 nM), (C) ARH3 (650 nM). Incubated samples were separated by RP-HPLC and observed at 254 nm as described in Materials and Methods. Purified peaks were confirmed using mass spectrometry. The assays were duplicated, and data represented a single separation. (●) ADP-ribose.

We next employed a dephosphorylation assay to monitor the release of ADPr from ADPr-1”-phosphate by SARS-CoV-2 Mac1. As expected, Mac1 released ADPr from the ADPr-1”-phosphate substate and the hydrolysis activity of Mac1 was similar to that of ARH3 (Figure 2 II). In contrast, SARS-CoV-2 Mac2 and Mac3 did not hydrolyze ADPr-1”-phosphate (data not shown).

Finally, we tested whether SARS-CoV-2 Mac1 hydrolyzes other ADPr O-linkage bond, i.e., ADPr-(serine)histone. ADPr-Ser was catalyzed by PARP-dependent ADP-ribosylation in the presence of HPF1 (28, 29).To generate ADPr-Ser, we used histone H3 peptide fragment (aa 1-21) as described in Materials and Methods, and serine mono-ADP-ribosylation was observed at serine 10 of histone peptide. As a control, we also confirmed that ARH3 cleaved ADPr-Ser (Figure 3D). Previous research showed that only ARH3 cleaved ADPr-Ser among various tested enzymes including macrodomain family (PARG, TARG1, MACROD1, and MACROD2) and ARH family (ARH1, ARH2, and ARH3) (30). We found that SARS-CoV-2 Mac1 was not able to catalyze removal of ADPr from the histone peptide (Figure 3C).

**Figure 3:**
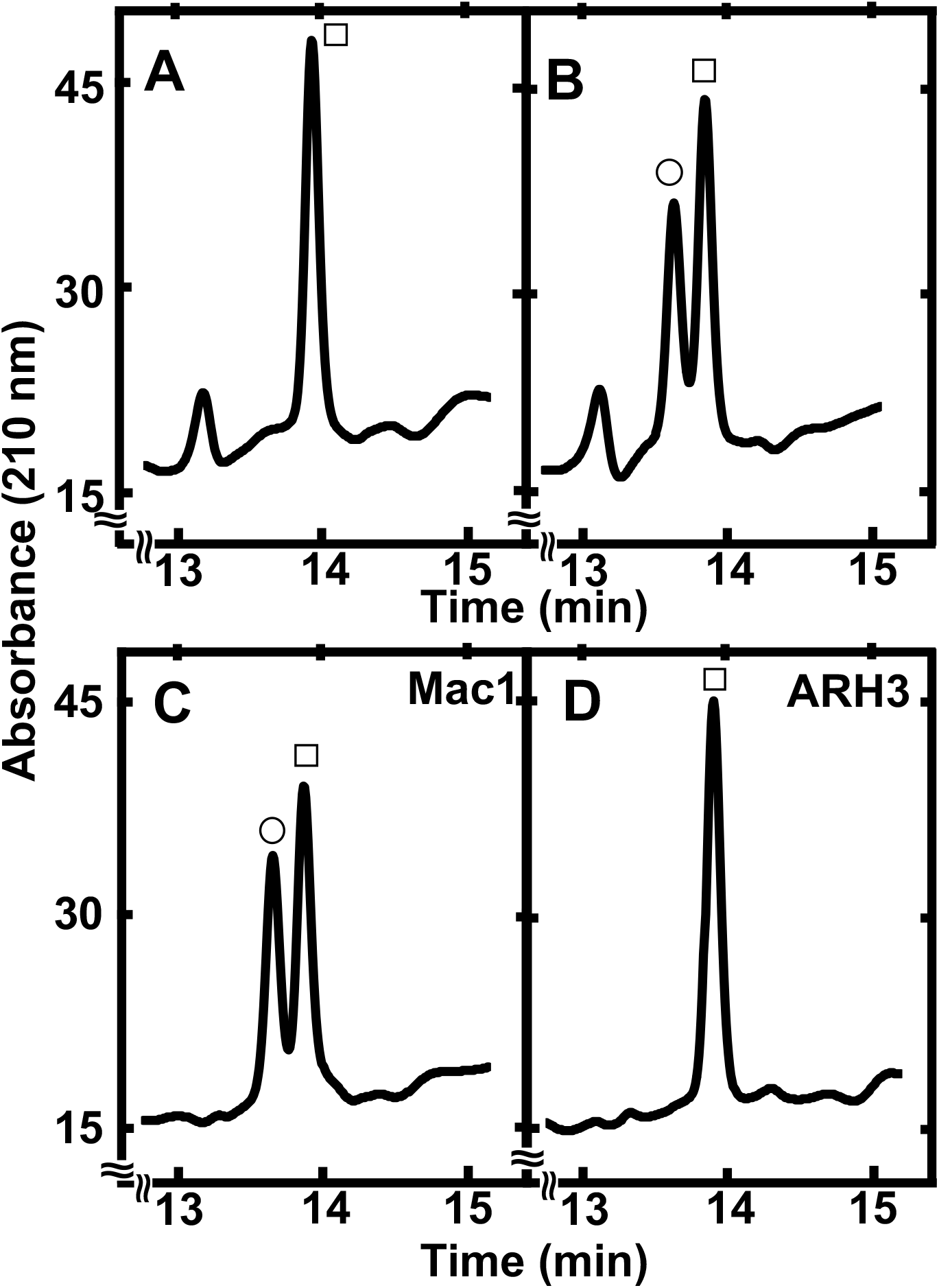
Hydrolysis activity of the ADP-ribose-serine by SARS-CoV-2 Mac1. Histone ADPr-Ser was modified in 20 μl of reaction mixture (RM) as described in material and method. The SARS-CoV-2 Mac1 (3 μM) and ARH3 (1 μM) were incubated with RM at 30°C, 30 min. Incubated samples was diluted with water containing 0.05% (v/v) trifluoracetic acid for a 200 μl total volume and analyzed by RP-HPLC detected 210 nm UV absorbance. (A) histone H3 peptide, (B) incubated ADPr-Ser, (C) SARS-CoV-2 Mac1, (C) ARH3. (☐) unmodified histone H3 peptide, (◯) ADP-ribosylated peptide.

### SARS-CoV2 Mac1 hydrolyzed α-NAD^+^ and *O*-Acetyl-ADPr without Mg^2+^

ARH3 was reported to hydrolyze ADP-ribose linked to O- (e.g., *O*AADPr) and N- (e.g., α-NAD^+^) functional groups (31, 32). In the catalyzing reaction, ARH3 was stimulated by Mg^2+^ and the reaction was enhanced by dithiothreitol (DTT) (31, 33). To determine if there was an effect of Mg^2+^ on the Mac1 reaction, reaction mixtures of Mac1 with and without Mg^2+^ were incubated with α-NAD^+^ or *O*AADPr. In contrast to the ARH3-catalyzed reaction, Mac1 reaction was not stimulated by Mg^2+^ (Figure 4 I-II). Similar activities to Mac1 bacterial Af1521 and human C6orf130 activities were not stimulated by Mg^2+^ (Figure 4I-II).

**Figure 4:**
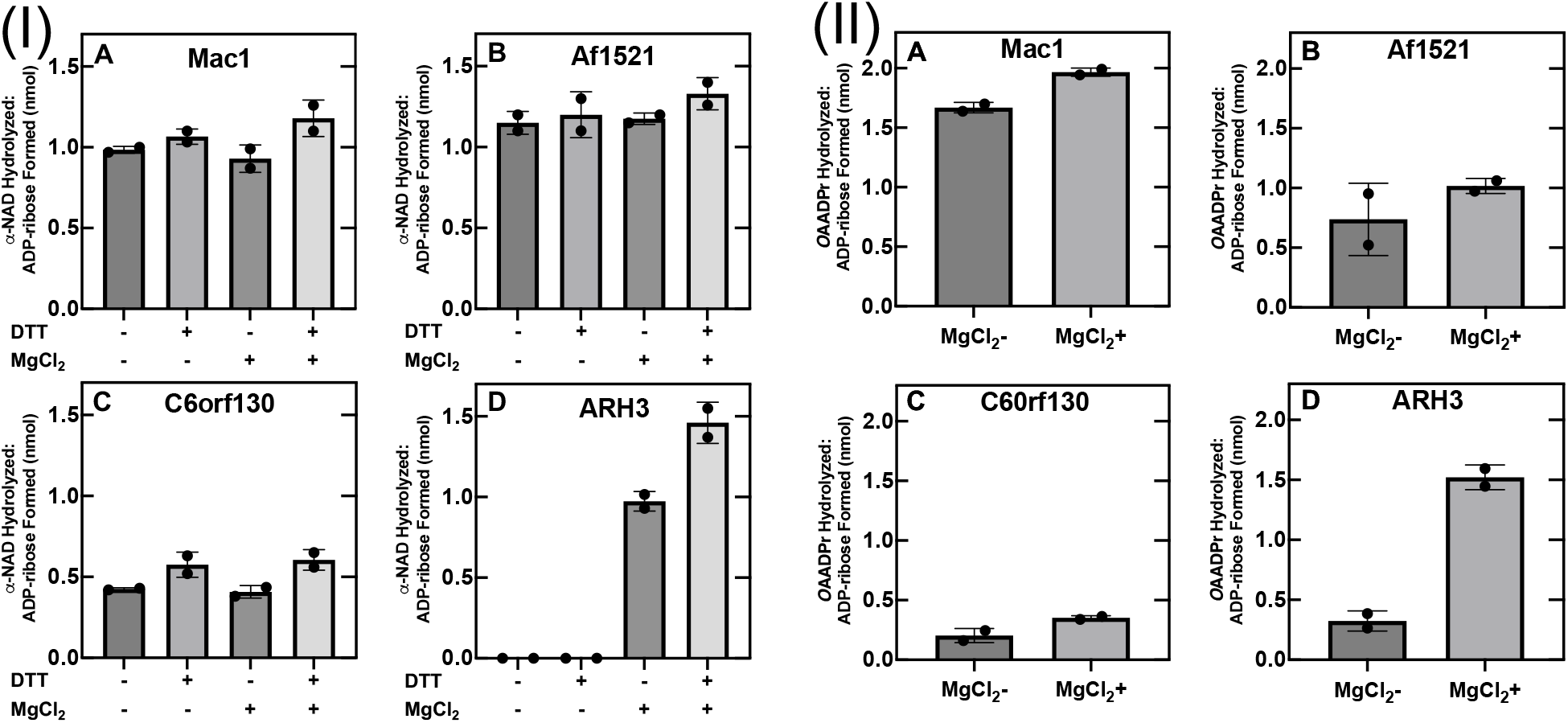
Characterization of α-NAD^+^ and *O*AADPr hydrolysis by Mac1, Af1521, C6orf130, and ARH3. (I) Effect of Mg^2+^ and DTT on *α*-NAD^+^ hydrolase activity. (A) The SARS-CoV-2 Mac1 (45 pmols), (B) *Archaeoglobus fulgidus* Af1521 (5 pmols), (C) human C6orf130 (15 pmols), and (D) human ARH3 (0.45 pmol) were assayed with *α*-NAD^+^ (20μM), 50 mM potassium phosphate (pH 7.0), with or without 10 mM MgCl_2_ and/or 5 mM DTT, and 22.5 μM ovalbumin, in 200 μl of reaction mix for 1h at 37°C before separation by RP-HPLC as described in Materials and Methods. (II) Requirement of Mg^2+^ for *O*AADPr hydrolase. (A) the SARS-CoV-2 Mac1 (1.5 pmol), (B) *Archaeoglobus fulgidus* Af1521 (1.5 pmol), (C) human C6orf130 (1.5 pmol), and (D) human ARH3 (1.5 pmol) were incubated with *O*AADPr (20μM), 50 mM potassium phosphate (pH 7.0), presence or absence of 10 mM MgCl_2_, and 22.5 μM ovalbumin, in 200 μl of reaction mix for at 25°C for 1h before separation by RP-HPLC as described in Materials and Methods. All assays were performed in duplicate assays. Data are shown as mean ± SD. Results were similar in both experiments.

## DISCUSSION

Severe acute respiratory syndrome coronavirus 2 (SARS-CoV-2) infections are associated with an acute inflammatory response with increased levels of multiple pro-inflammatory cytokines result in an uncontrollable cytokine storm (34).

Macrodomains are conserved domain throughout all kingdoms of life. Thus, SARS-CoV-2 genome and its Mac1 shared >80% similarity with SARS-CoV and 72% similarity with SARS-CoV Mac1, respectively (24). Macrodomain Mac1 of SARS-CoV-2 is essential for viral replication and pathogenesis. The macrodomains have enzymatic ability to dephosphorylate ADP-ribose-1”-phosphate (a byproduct of RNA splicing reactions), which was first found in yeast (35) and in viruses (15, 36). In this study, SARS-CoV-2 Mac1 hydrolyzed ADP-ribose 1” phosphate (Figure 2, II, B). This finding may suggest the de-ADP-ribosylation by SARS-CoV-2 is specific at the catalytic site residing in ADP-ribose moiety at 1” position, which is similar to position of ADP-ribose-1”-phosphate. Interestingly, recent studies on antiviral activity have focused on signal-generating enzyme that processed NAD^+^ to generate ADP-ribose derivatives including ADP-ribose-1”-phosphate as a second messenger; this toxic metabolite can cause cell death (37). Therefore, to gain better understanding about involvement of ADP-ribose-1”-phosphate, observing other roles of SARS-CoV-2 Mac1 on other substrates would be an interesting field to study.

NAD^+^ is a mediator of both anti-viral and anti-inflammatory responses, which is involved in protein-protein interactions and served as a marker of energy (38, 39), play a crucial role in fueling enzymatic activity to support host immune regulation and responses (40). Lower concentrations of NAD^+^ were found in human peripheral blood leukocytes infected with HIV-1 *in vitro*, human fibroblast herpes simplex virus 1, and skeletal muscle of patients with HIV-1 and hepatitis C (41–43). Parallel to these, there is increasing of evidence showing decreased concentration of nicotinamide mononucleotide (NMN), an NAD^+^ precursor, in patients with severe COVID-19 symptoms compared to healthy subjects (44). Our study found that SARS-CoV-2 Mac1 cleaved α-NAD^+^, generating ADP ribose and nicotinamide, but not β-NAD^+^ (Figure 1, I). This substrate preference is consistent with Mac1 being stereospecific at the C-1” bond. In addition, there was *in vitro* investigation, using DBT, 17Cl-1, HEK293T, and HeLa cell lines infected with SARS-CoV-2 on the changes in some gene expressions including PARP9, PARP12, PARP14, and nicotinamide phosphoribosyltransferase (NAMPT), which consumed NAD^+^ for their synthesis, which was consistent with results seen in lung tissue and bronchoalveolar lavage fluid (BALF) of COVID-19 patients (45). Given the facts above, depletion of NAD^+^ found in COVID-19 patients may have resulted from cleavage of α-NAD^+^ by SARS-CoV-2 macrodomain Nsp3 observed specifically by Mac1 but not Mac2 and Mac3 (data not shown) during infection and other effects probably due to the high utilization of β-NAD^+^ by enzymes including sirtuin, CD38 and PARPs during infection (46). To fight against viral infection maintaining cellular β-NAD^+^ levels of host cells, one of the host defense strategies suggests to disable the contribution of viral NAD^+^ consumption for viral replication, and promote NAD^+^-consuming domains of anti-viral enzymes, e.g., sirtuin domains (40).

Sirtuins are the NAD^+^-consuming enzymes that involved in immune responses (40). Particularly, silent information regulator 2 (Sir2) family of enzymes are involved in cellular processes included histone deacetylation, chromatin stability, aging, and gene silencing (47). In the presence NAD^+^, a marker of energy availability and an acceptor to catalyze the diacetylation of protein, Sir2 splits groups from protein and yielding *O*-acetyl-ADP-ribose and nicotinamide (48). *O*-acetyl-ADP-ribose plays an important role in stabilization of chromatin remodeling and assembly of Sir2-histone complexes (49, 50), cellular metabolism, and redox signaling (51, 52). There are reports showing that macrodomain-containing proteins including human MacroD1, human MacroD2, *E.Coli* YmdB, *Staphylococcus aureus* sirtuin-linked SAV0325, human C6orf130 possessed *O*-acetyl-ADP-ribose hydrolase activity, catalyzing the deacetylation of *O*-acetyl-ADP-ribose to produce ADP-ribose and acetyl as the products (27, 53). Of note, in our present study, SARS-CoV-2 Mac1 of Nsp3 cleaved *O*-acetyl-ADP-ribose to generate ADP-ribose as the product, whereas Mac2 and Mac3 did not (data not shown)., suggesting that SARS-CoV-2 containing Nsp3 might be involved in chromatin stabilization and cellular metabolism to increase its virulence and suppress host cellular anti-viral activity against COVID-19 infection.

In this report, we used ARH1 and ARH3 as positive controls of enzymatic activity assays, confirming previous findings that human ARH3 hydrolyzed ADP-ribose joined to O- and N-functional groups (30–33, 54). As expected, human ARH3 hydrolyzed ADP-ribose-serine and *O*-acetyl-ADP-ribose. Of interest, our data showed that ARH3 cleaved α-ADP-ribose-arginine bond, although ARH3 has approximately 250 times less active than ARH1 (Figure 1E; Table 1). Using this system, we found that SARS-CoV-2 Mac1, but not Mac2 and Mac3, specifically cleaved various substates, with ADP-ribose attached to O*-* and N*-*linked functional groups, α-NAD^+^, *O*-acetyl-ADP-ribose, and ADP-ribose-1”-phosphate, but did not hydrolyze α-ADP-ribose-arginine and ADP-ribose-(serine)histone H3, as did ARH1 and ARH3 (Figure 1, IIC; Figure 2, IB; Figure 2, IIB; Figure 3). ARHs catalyzes linkages of ADP-ribose and residues including arginine and serine in a Mg^2+^-dependent and, in some species, in a thiol-dependent manner (33, 55–57). In contrast, similar to other macrodomains from bacteria Af1521 and mammalian C6orf130, SARS-CoV-2 hydrolyzed substrates both O- and N-linked to ADP-ribose in the absence of Mg^2+^ (Figure 4).

**Table 1:** Hydrolase activities of macrodomain-containing proteins and ARHs families.

| Proteins | Substrates |  |  |  |  |  |
| --- | --- | --- | --- | --- | --- | --- |
| | α-NAD | β-NAD | ADPr-arg | ADPr-1" P | OAADPr | ADPr-Ser |
| **ARHs** |  |  |  |  |  |  |
| ARH1 | yes<sup>a</sup> | no<sup>a</sup> | yes<sup>a*</sup> | yes<sup>*</sup> | yes<sup>a</sup> | ----- |
| ARH3 | yes<sup>*</sup> | no<sup>*</sup> | yes<sup>*</sup> | yes<sup>*</sup> | yes<sup>*</sup> | yes<sup>b</sup> |
| **Macrodomains** |  |  |  |  |  |  |
| Mac1 | yes<sup>*</sup> | no<sup>*</sup> | no<sup>*</sup> | yes<sup>*</sup> | yes<sup>*</sup> | No<sup>*</sup> |
| Mac2 | no<sup>*</sup> | no<sup>*</sup> | no<sup>*</sup> | no<sup>*</sup> | no<sup>*</sup> | ----- |
| Mac3 | no<sup>*</sup> | no<sup>*</sup> | no<sup>*</sup> | no<sup>*</sup> | no<sup>*</sup> | ----- |
| C6orf130 | yes<sup>*</sup> | no<sup>a</sup> | no<sup>a</sup> | ----- | yes<sup>*</sup> | ----- |
| Af1521 | yes<sup>*</sup> | no<sup>a</sup> | no<sup>a</sup> | ----- | yes<sup>*</sup> | ----- |

Mac1, SARS-CoV-2 Mac1; Mac2, SARS-CoV-2 Mac2; Mac3, SARS-CoV-2 Mac3. ADPr-arg, ADP-ribose-arginine; ADPr-1" P, ADP-ribose-1"-phosphate; OAADPr, O-Acetyl-ADP-ribose; ADPr-Ser, ADP-ribose-serine; (\*) indicates this manuscript. "yes or no" indicates activity has been reported for the respective substrates. Dashes (---) represent activities which have not been reported. a<sup>28</sup>, b<sup>29</sup>.

In summary, SARS-CoV-2 Nsp3 Mac1 can hydrolyze ADP-ribosylated substrates involved in immune responses and DNA repair via α-NAD^+^, ADPr-1”-phosphate, and *O*-acetyl-ADP-ribose. In particular, SARS-CoV-2 Mac1 appears to function in viral replication and pathogenesis and to exert antiviral activity by affecting host-mediated ADP-ribosylation. Thus, inhibiting activities of Nsp3 Mac1 or stabilizing their substrates may be critical to blocking SARS-CoV-2 activity.

## MATERIALS AND METHODS

### Reagents

α-NAD, β-NAD, cyclic adenosine diphosphate-ribose (C7344), L-arginine, dithiothreitol (DTT), ovalbumin, HPLC water with 0.05% (v/v) trifluoracetic acid (TFA) or with 0.05% (v/v) acetonitrile, and histone H3 peptide (aa 1-21) were purchased from Sigma Aldrich. Cholera Toxin A subunit (CTA) was purchased from List Biological Laboratories (CA, USA). *O*-acetyl-ADP-ribose was obtained from Toronto Research Chemicals (Toronto, Canada). Human PARP1 enzyme and human HPF1 protein were purchased from Tulip Biolabs (West Point, PA, USA).

### Protein expression and purification

Recombinant human ARH1, human ARH3, human C6orf130, and *Archaeoglobus fulgidus* Af1521were prepared as described previously (58, 59). SARS-CoV-2 non-structural proteins (Nsp) 3 macrodomains (Mac) 1, 2, and 3 were cloned into a pETM33 expression vector with His_6_ and GST tags. For the expression of protein used for enzymatic assays, the plasmid DNAs were transformed into the *E. coli* Rosetta strain BL21(DE3). A single colony of plasmid harboring cells was transferred into 25 ml LB medium (Miller; MD, USA) supplemented with kanamycin (50 μg/ml) and cultured overnight at 37°C, with shaking at 225 rpm. Large scale culture was conducted by using 10 mL of the starter cultures, transferred to, and cultured in a shaker flask containing 250 mL of LB medium at 37°C for 12 h. The proteins were induced with 1 mM isopropyl-β-D-thiogalactopyranoside (IPTG) at 37°C for 6 h. Cells were harvested, and pellets were collected at 4,500 rpm for 10 min and frozen at −80°C. Frozen cells were thawed on ice and used for protein purification with nickel-nitrilotriacetic acid (Ni-NTA) Fast Start column as recommended by the manufacturer (Qiagen, Germany). The bound proteins were eluted from the column with elution buffer containing 50 mM NaH_2_PO_4_, 300 mM NaCl and 250 mM imidazole at pH 8.0, followed by dialysis using phosphate buffered saline (PBS) overnight at 4°C.

### Generation and analysis of ADP-ribose-arginine (ADPr-arg)

ADPr-arg was generated using bacterial toxin ADP-ribosyltransferase cholera toxin A subunit (CTA) in a reaction mixture of CTA (0.2μg/μl), 30 mM DTT, 1 mM β-NAD, and 50 mM L-arginine with 0.17 μg/μL ovalbumin in 20 mM potassium phosphate (pH 7.0, volume 300 μL), which was incubated at 30 °C for 3 h. The incubated reaction mixtures (200 μl) were injected into HPLC. The products were separated on a silica-based Discovery Bio Wide Pore C18 column (25cm x 4.6mm, 5 μm particle size; Supelco, Bellefonte, PA, USA) by gradient elution with flow rate of 1 mL/min, equilibrated with 100% buffer A (0.05 (v/v) trifluoroacetic acid (TFA) in HPLC grade water) from 0 to 8 min, followed by a linear gradient to 100% buffer B (0.05 (v/v) TFA in acetonitrile) from 8 to 18 min, and detected under UV absorbance at 254 nm. ADPr-arg was collected and concentrated by vacuum evaporator, and product was confirmed by mass spectrometry analysis.

### Enzymatic hydrolysis of ADP-ribose-arginine, α-NAD^+^, and β-NAD^+^

Incubated samples of ADPr-arg (35 μM, 7 nmols), α-NAD^+^ (20 μM), or β-NAD^+^ (20 μM), and purified recombinant proteins of SARS-CoV2 Nsp3 (i.e., Mac1, Mac2, and Mac3), human ARHs, *Archaeoglobus fulgidus* Af1521, or human C6orf130, were incubated in 50 mM potassium phosphate buffer (pH 7.0), 10 mM MgCl_2_, 5 mM DTT, and 22.5 μM ovalbumin (total volume 200 μl) at 37°C. The products were then separated by HPLC and confirmed by mass spectrometry.

### Hydrolysis of *O*-Acetyl-ADPr Catalyzed by macrodomain proteins and ARH3

*O*AADPr substrate and purified recombinant SARS-CoV-2 Nsp3 (i.e. Mac1, Mac2, and Mac3), human ARH3, bacterial Af1521, or human C6orf130 were incubated in 50 mM potassium phosphate buffer (pH 7.0), 10 mM MgCl_2_, and 22.5 μM ovalbumin (total volume 200 μl) at 25°C for 1 h. Products were separated and analyzed by RP-HPLC, on silica-based Discovery Bio Wide Pore C18 column (25cm x 4.6mm, 5 μm particle size; Supelco, Bellefonte, PA, USA), with elution gradient 5-15 min of 90-100% solvent A (0.05% (v/v) TFA in water) and 0-10% solvent B (0.05% (v/v) TFA in acetonitrile) and confirmed by mass spectrometry.

### Isolation of ADP-ribose-1”-phosphate (ADPr-1”-phosphate)

A portion of 0.1 mM of cADP-ribose was fractionated using Supelcosil LC-18-T (Supelco) HPLC column (15 cm x 4.6 mm, 3 μm) equilibrated with mobile phases A and B as described previously (32). Briefly, 200 μl of incubated samples were separated with a gradient mobile phase with buffer A (0.1M sodium phosphate/4 mM tetrabutylammonium hydrogen sulfate, pH 6.0) for 2.5 min, followed by a mobile phase with 30% buffer B (buffer A: methanol, pH 7.2; ratio 70:30) from 2.5-5.0 min, and continued to a gradient up to 60% buffer B from 5.0 to 10.0 min. The flow rate was maintained at 1 ml/ min. Fractions of cADP-ribose was eluted at 3 min, ADP-ribose was at 8 min and ADPr-1”-phosphate, which is a byproduct, was eluted at 10 min. The eluted peaks were confirmed by using mass spectrometry.

### Hydrolysis of ADPr-1”-phosphate

Assays containing 6 μM of purified ADP-ribose-1”-phosphate and enzymes were incubated overnight in a 50 mM Tris-HCl (pH 7.5) buffer containing 10 mM MgCl_2_, 5 mM DTT, and 22.5 μM ovalbumin in a total volume of 200 μl at 37°C. Substrate and products were separated by using HPLC as described above.

### Identification of ADP-ribose-serine (ADPr-Ser) in histone H3 by HPLC-mass spectrometry

ADPr-Ser substrate was generated by a reaction mixture of PARP1, HPF1, histone H3 (aa 1-21) peptide, and NAD^+^ as described (30). Briefly, recombinant histone H3 (aa 1-21) peptide (0.25 nmol, 1.25 μM) was ADP-ribosylated by 1μM PARP1 and 1μM HPF1 in a reaction buffer (total reaction volume, 20 μl) containing 50 μM NAD^+^, 50 mM Tris-HCl pH 8.0, 10 mM MgCl_2_, 100 mM NaCl at 30°C for 30 min. The reaction was stopped by addition of 1 μM of PARP inhibitor Olaparib.

ADPr-Ser-histone H3 peptide was digested into smaller fragments by endopeptidase prior to the analysis. Briefly, after drying the reaction mixture with a Savant SpeedVac (Thermo Fisher Scientific, MA, USA), peptides were resuspended in protease reaction buffer (100 mM Tris, 1mM diethylenetriamine pentaacetate (DTPA), pH 8.5) and digested with Lys-C endopeptidase (Mass Spectrometry Grade, Wako Chemicals, Japan) for 18 h at 37 °C with a ratio of Lys-C:histone H3 peptide of 1:25. The reaction was stopped by 10% TFA and peptides were analyzed by Agilent 6530C Quadrupole-Time of Flight (QTOF) LC/MS (Agilent Technologies, DE, USA). Digested peptides were separated on ZORBAX 300SB-C18 MicroBore column (1.0 x 50 mm, 3.5mm, Agilent Technologies, DE, USA) with a linear gradient of acetonitrile from 0 to 50% at 1% per min with a flow rate of 20 ml/min. Eluents from the column was mixed in a mixing tee with glacial acetic acid at 15 ml/min flow rate prior to the electrospray nebulizer to exchange the bound TFA (60). Positive electrospray ionization spectra were obtained in the mass range of 100 to 2500 m/z. The drying gas temperature was 350 °C with a flow of 10 L/min and a nebulizer pressure of 30 psi. Voltages of the capillary and fragmentor were 3500 V and 235 V, respectively. MS/MS fragmentation used variable collision energies of 9 ~ 69 eV for 300 ~ 1500 m/z with a data collection mass range of 50 to 2000 m/z. MS and MS/MS spectra were analyzed using Agilent software, MassHunter version B.07. To identify ADP-ribosylation on serine residue of histone H3, MS/MS spectra were analyzed using a software platform, PEAKS Studio Xpro (Bioinfomatics Solutions Inc., Ontario, Canada). Predicted MS/MS spectra were generated by GPMAW ver. 12 (Lighthouse Data, Odense, Denmark) and matched to the experimentally obtained spectra.

### ADPr-Ser hydrolase activity assays

The ADP-Ser ribosylated peptide was used as a substrate and incubated with various enzymes, e.g., SARS-CoV-2 Mac1 (3 μM), human ARH3 (1μM), for 30 min at 30°C. Reaction mixtures were then subjected to RP-HPLC, using solvent A (0.05% TFA in water) and 0.05% TFA in acetonitrile (solvent B), with gradient 2-22 min of 0 to 35% solvent B. The purified peaks were identified by mass spectrometry.

### Identification of ADPr by HPLC-Mass Spectrometry

HPLC-mass spectrometry analysis was performed on an Agilent 6530C quadrupole-time of flight (QTOF) mass spectrometer with an Agilent 1200 series capillary system. To separate analytes, we used two different modes of separation, reverse-phase liquid chromatography (RPLC) and hydrophilic interaction liquid chromatography (HILIC). For the RPLC, the LC is equipped with a KINEX 2.6 μm Polar C18 (2.1 x 50 mm, 100 Å, Phenomenex, CA, USA) set to 30°C. The initial solvent was 10 mM ammonium acetate acid (solvent A), and analytes were eluted by a linear gradient of 2%/min acetonitrile (solvent B) from 0 to 60 % with a flow rate of 60 μl/min. For the HILIC, the LC is equipped with an InfinityLab Poroshell 120 HILIC-z (2.1 x 50 mm, 2.4 μm, Agilent Technologies, DE, USA). The initial solvent was 10 mM ammonium formate in 90 % acetonitrile (solvent B). The analytes were eluted by increasing at 4%/min 10 mM ammonium formate (solvent A) from 0 to 100% with a flow rate of 60 μl/ min. Positive electrospray ionization MS/MS spectra were obtained in the mass range of 50 ~ 1300 m/z with collision energy 15-25 eV. The drying gas temperature was 300°C with a drying gas flow of 8 L/min and a pressure of 25 psi. The capillary and fragmentor voltages were 3500 V and 120 V, respectively. Mass spectra were analyzed using Agilent software, MassHunter Qualitative Analysis Version B.07 (Agilent Technologies, CA, USA). Spectral matching of obtained MS/MS spectra was performed with spectral libraries Human Metabolome Database (HMDB) and METLIN Gen2.

## ACKNOWLEDGMENT

We thank Rodney L. Levine (Laboratory of Biochemistry; NHLBI; NH) for providing valuable suggestions. This research was funded by the Intramural Research Program of National Institutes of Health (NIH), the National Heart, Lung, and Blood Institute.

## Author contributions

C.C. and J.M. designed experiments. D.Y.L., J.K., and H.I.E. helped to design experiments. J.K. prepared recombinant proteins. C.C. and D.Y.L. performed experiments. C.C., J.M., D.Y.L., H.I.E. analyzed data. C.C. and J.M. wrote the manuscript. All authors reviewed the manuscript.

